# Whole-Slide ECM Imaging Reveals Dense Fibrous Matrix as a High-Risk Factor for Recurrence in Stage II Colon Cancer

**DOI:** 10.1101/2024.11.14.622985

**Authors:** C.J. Ravensbergen, V.S. Colaco, H. Putter, A.S.L.P. Crobach, J. Boonstra, J.H.F. Lindeman, R.A.E.M. Tollenaar, W.E. Mesker

**Author notes:** Correspondence to: Dr. Wilma Mesker, Department of Surgery, Leiden University Medical Center, Albinusdreef 2, P.O. Box 9600, 2300RC Leiden, the Netherlands. Mail.

## Abstract

**Introduction:** The extracellular matrix (ECM) supports tumor progression by influencing tumor cell migration and invasion. This study examines the link between peritumoral ECM morphology and five-year recurrence risk in TNM Stage II colon cancer, using quantitative whole-slide ECM imaging. We hypothesize that loose ECM regions are associated with increased recurrence risk due to enhanced tumor budding (TB) or poorly differentiated clusters (PDC).

**Methods:** In a case-control study of 100 TNM Stage II colon cancer patients (25 with recurrence and 75 controls matched by lymph node sampling and tumor extent), Picrosirius red-stained sections were imaged to quantify ten ECM parameters across 798 regions of interest (ROIs). Conditional logistic regression assessed associations between ECM morphologies, TB, PDCs, and recurrence.

**Results:** Unsupervised clustering identified three ECM morphologies: dense fibrous, loose sparse, and complex tortuous. Dense fibrous ECM correlated strongly with recurrence (aOR 9.43, 95% CI 3.29-29.30, *p* < 0.001), while loose sparse and complex tortuous ECMs were associated with reduced recurrence risk (aOR 0.33, 95% CI 0.11-0.91, *p* = 0.040, and aOR 0.14, 95% CI 0.02-0.53, *p* = 0.012, respectively). TB was highest in loose sparse ECM (mean 9.3), and PDCs were highest in dense fibrous ECM (mean 5.5).

**Discussion:** Our findings suggest that ECM morphology, particularly dense fibrous ECM, predicts recurrence in Stage II colon cancer, highlighting ECM profiling as a promising tool for patient stratification beyond traditional staging.

## Introduction

The tumor-stroma ratio (TSR) is closely related to tumor aggressiveness, with high stromal content being a well-established indicator of poor prognosis in solid tumors [1-4]. As a key component of tumor stroma, the extracellular matrix (ECM) provides structural support and helps drive tumor migration [5]. During migration, tumor cells reciprocally interact with the ECM, altering its structure to enhance invasion [6]. This dynamic interaction suggests that local ECM morphology directly shapes tumor migration patterns. For instance, alignment of the fibrous ECM perpendicular to the tumor border has been shown to facilitate collective migration of tumor cells [7-10]. Additionally, we recently demonstrated that a loose collagen matrix is associated with single-cell migration and lymph node metastasis in early-stage colon cancer [11].

Histological markers of tumor migration, such as tumor budding, poorly differentiated clusters, and border configuration, likely mirror the underlying ECM morphology. Tumor budding—characterized by single cells or small clusters of cells at the invasive front—has emerged as a marker of aggressive tumor behavior and poor prognosis in TNM Stage II colon cancer [12-16]. The budding process is closely tied to epithelial-mesenchymal transition (EMT), where budding cells lose cell-adhesion molecules like Ep-CAM and upregulate mesenchymal markers like vimentin [15, 17-21]. Stage II colon cancer comprises a heterogeneous group, with up to 15% of patients experiencing disease recurrence within five years after curative surgical resection [14, 16, 22]. For those at high risk of recurrence, adjuvant systemic treatment may improve outcomes. While tumor budding enhances recurrence prediction beyond traditional TNM staging, variability in assessment and the localized focus of hot-spot analysis may overlook broader tumor heterogeneity [23, 24].

Unlike the small clusters of budding cells that can be difficult to detect, the ECM at the tumor border is often abundant and spans across the peritumoral stroma, offering an accessible marker for analysis. Given the ECM’s role in driving tumor migration, this study aims to explore how peritumoral ECM morphology relates to five-year recurrence risk in TNM Stage II colon cancer. We hypothesize that regions with loose ECM morphology are linked to higher tumor recurrence, possibly due to their association with increased tumor budding. Using the fluorescent properties of Picrosirius red histochemical staining, we introduce a practical method for quantitative whole-slide ECM characterization in histological tissue sections.

## Methods

### Study Cohort and Case-Control Matching

This study employed a case-control design. The study cohort comprised patients with pathologic TNM (eighth edition) stage II colon carcinoma who underwent curative surgical resection at Leiden University Medical Center between 2003 and 2018. All patients were therapy-naïve before surgery. Clinical data and pathology reports were retrospectively collected from the electronic patient files. Patients were eligible for inclusion if they had complete follow-up data for a minimum of five years from the date of surgery or until recurrence occurred. Cases were patients who experienced locoregional or distant disease recurrence within five years from the date of surgery. Controls were patients who did not experience disease recurrence within a five-year follow-up period after surgery. Controls were matched to cases without replacement based on two major clinicopathologic risk factors for recurrence in stage II disease, as defined by ASCO/ESMO: tumor extent (pT3 or pT4 status) and the number of lymph nodes sampled [14, 16]. For the number of lymph nodes sampled, a control was selected if their value fell within a margin of ±5 nodes from the value of the case. For each case, three unique controls (1:3 ratio) were randomly selected from the respective matched pool members. Formalin-fixed paraffin-embedded (FFPE) resection tissue of the most invasive part of the tumor (diagnostic T status slide) was obtained for ECM imaging. All tissue samples were coded and handled according to ethical standards (‘Code for Proper Secondary Use of Human Tissue’, Dutch Federation of Medical Scientific Societies). This study was approved by the Medical Ethics Committee of the LUMC (ID G21.093).

### Tissue Processing, Histochemical Staining and Digitization

Paraffin-embedded tumor tissues were sectioned into four µm slides and underwent sequential processing including deparaffinization with xylene, rehydration through ethanol gradients, histological staining, air drying at room temperature, and mounting with a xylene-based mounting medium (Permount; Thermo Fisher Scientific Inc., Waltham, Massachusetts, USA). In this study, one tissue block containing the most invasive part of the tumor, used to determine the T-status, was used to prepare serial tissue sections. These serial tissue sections were subjected to various histochemical stains. A saffron-enhanced hematoxylin & eosin (HES) staining was performed using a routine H&E protocol, followed by an additional incubation step in a saturated alcoholic saffron solution. Sections were treated with Picrosirius red (0.1% Direct Red 80 dye in saturated picric acid; Sigma-Aldrich, Merck, Darmstadt, Germany) for 60 minutes, then rinsed in two changes of 0.5% glacial acetic acid (Avantor, Inc., Radnor Township, Pennsylvania, USA). Herovici’s staining kit was applied to distinguish between reticulin and mature collagen, according to manufacturer instructions (ScyTek Laboratories, Inc., Logan, Utah, USA). Elastic fibers were stained using the Taenzer-Unna orcein method, where tissue sections were incubated in a 1% orcein solution (Sigma-Aldrich, Merck, Darmstadt, Germany) in 1% hydrochloric acid and 70% alcohol at room temperature for 60 minutes, followed by a five-second differentiation step in 1% hydrochloric acid in 70% alcohol. Finally, to visualize glycosaminoglycans (GAGs), sections were stained with alcian blue pH 2.5 solution (Sigma-Aldrich, Merck, Darmstadt, Germany) for 30 minutes and counterstained with nuclear fast red solution (Sigma-Aldrich, Merck, Darmstadt, Germany) for 5 minutes. Movat’s Pentachrome was performed through sequential incubations with Alcian Blue (pH 2.5), Weigert’s Hematoxylin, Crocein Scarlet Acid Fuchsin, and Alcoholic Saffron (Sigma-Aldrich, Merck, Darmstadt, Germany). Stained tissue sections were digitized in brightfield mode using a Pannoramic 250 slide scanner (3DHISTECH, Budapest, Hungary) at ×20 magnification (0.39 µm per pixel) and saved as proprietary .mrxs files.

### Fluorescent Whole-Slide Imaging of Picrosirius Red-Stained Tissue Sections

Picrosirius red stained tissue sections were digitized in fluorescent mode using a Pannoramic 250 slide scanner (3DHISTECH, Budapest, Hungary) at ×20 magnification (0.39 µm per pixel) and saved as proprietary .mrxs files. Tissue autofluorescence was detected using a FITC filter cube (excitation 460-488nm, emission 502-547) at 800ms exposure time. The emission of the fibrous ECM stained by Sirius red was detected using a TRITC filter cube (excitation 532-554nm, emission 573-613) at 44ms exposure time.

### ECM Image Analysis Pipeline

For quantitative ECM analysis, regions of interest (ROIs) in the peritumoral stromal areas along the invasive border were identified and annotated in multichannel fluorescent whole-slide images of Picrosirius red-stained tissues. The ROIs corresponded to those typically used for histologic scoring of peritumoral budding. Each ROI was annotated using a circular area of 0.785 mm^2^, which represents the microscopic field of view at 200x magnification. Depending on tissue availability, up to ten ROIs were annotated per tumor section, with a minimum of five ROIs. Regions with necrosis and dense cellular infiltrate were avoided. We measured ten quantitative ECM parameters to analyze the structural characteristics within each ROI (*Supplementary Figure S1*). Briefly, the anisotropy index assessed the directional alignment of fibers, while the dominant direction indicated their primary orientation. Branching density and intersection density quantified the complexity and connectivity of the ECM network. Average fiber length provided the typical length of ECM fibers. Fractal dimension measured structural complexity, and lacunarity captured the distribution of gaps and voids, indicating heterogeneity. Fiber bundle density determined the packing density of fibers, and tortuosity quantified the winding nature of fiber paths. Lastly, the matrix area-perimeter ratio assessed the compactness of ECM structures, with higher values indicating more compact shapes. The parameters were obtained using Matrix ORientation and Texture EXplorer (MORTEX), an in-house developed ImageJ (version 1.54j) macro that serves as a wrapper for dependencies such as Bio-formats, AnalyzeSkeleton, OrientationJ, and ComsystanJ. [25-29]. The source code and instruction manual of the pipeline are publicly available at https://www.github.com/cjravensbergen/MORTEX. MORTEX is publicly available under the terms of the GNU General Public License (GPL) v3.

### Picrosirius Red Polarization Microscopy

Picrosirius red-stained sections were imaged on a Leica DM6 B upright microscope with a DCF495 camera and 20x/0.7 N.A. oil-immersion objective (Leica Microsystems, Mannheim, Germany), using polarization contrast microscopy to detect birefringent collagen fibers. Under polarized light, collagen displayed red, orange, yellow, and green hues based on fiber orientation and density. Optimal exposure settings were adjusted to capture vivid birefringence without intensity saturation, with consistent white balance, gain, and exposure applied across images. The images were saved as 16-bit RGB .lif files.

### Immunohistochemistry

Tissue sections were stained for cytokeratin using single immunohistochemistry. To improve antigen accessibility, the slides underwent heat-induced epitope retrieval in a 10 mM citrate buffer (pH 6.0). Endogenous peroxidase activity was blocked by incubating the slides with 1% hydrogen peroxide (H_2_O_2_) for 20 minutes at room temperature. The slides were then incubated overnight at room temperature with a monoclonal mouse anti-human cytokeratin primary antibody (clones AE1/AE3, M3515, Dako, Glostrup, Denmark) at a 1:300 dilution in phosphate-buffered saline with 1% bovine serum albumin. Visualization was performed using the EnVision+ detection system (Dako, Glostrup, Denmark) with 3,3’-diaminobenzidine (DAB) as the chromogenic substrate. Finally, the slides were counterstained with Mayer’s hematoxylin (Merck Millipore, Amsterdam, the Netherlands) for 30 seconds, rinsed with running tap water, air-dried, and mounted using a synthetic resin medium.

### Tumor-Stroma Ratio, Tumor Budding, and Poorly Differentiated Clusters

The tumor-stroma ratio (TSR) was estimated on HES-stained tissue sections following a previous protocol [30]. A hotspot (one per tumor) circular annotation of 3.14 mm^2^, corresponding to a 100x magnification microscopic field of view, was placed in the tumor region with the highest amount of stroma. The percentage of stroma within the annotation was estimated in increments of 10%. Tumors were categorized as stroma-high (>50%) or stroma-low (≤50%). Tumor budding (TB) counts were estimated on tissue sections stained for cytokeratin. A tumor bud was defined as a single or small cluster of up to four tumor cells. TB was scored per the recommendations of the International Tumor Budding Consensus Conference (ITBCC) 2016 [12]. In our study, TB scoring was limited to the estimation of peritumoral bud counts. Intratumoral budding was not included in the analysis. TB counts were estimated in all ROIs used for ECM analysis and reported as continuous counts per ROI. For logistic regression analysis, TB counts were categorized into low-grade (Bd1, 0-4 buds), intermediate (Bd2, 5-9 buds), or high-grade budding (Bd3, ≥10 buds) groups. As an extension to TB, poorly differentiated clusters (PDC) were defined as small clusters of 5 or more tumor cells without glandular differentiation. PDCs were scored according to current recommendations [31]. For logistic regression analysis, PDC counts were categorized into low-grade (G1, 0-4 PDCs), intermediate-grade (G2, 5-9 PDCs), or high-grade budding (G3, ≥10 PDCs) groups.

### Statistical Analysis

R (v4.4.2; https://www.r-project.org/) was used for statistical analysis and data visualization. To evaluate the case-control matching process, the balance of matching variables and covariates between cases and controls was assessed using descriptive statistics and statistical tests. Variable distribution was assessed using the Shapiro-Wilk test, followed by parametric or non-parametric testing. Categorical variables were analyzed with Fisher’s exact test or Chi-squared test. Continuous variables were analyzed with unpaired t-test or Mann-Whitney U test and reported as median value and interquartile range. To reduce the dimensionality of the dataset and identify the most significant patterns, Principal Component Analysis (PCA) was applied to the dataset of ECM features across all ROIs. The number of retained principal components (PCs) was based on the minimal number of PCs needed to explain more than 80% of the variance in the data. After PCA, unsupervised hierarchical clustering was applied to the retained PCs using Ward’s method and the Euclidean distance. Cluster number selection was guided by average silhouette, visual inspection of clustering, and biological relevance. The resultant cluster assignments were visualized in a dot plot without further data transformation. Mean TB and PDC counts including 95% confidence interval (CI) were reported. To analyze the association between ECM morphology and recurrence, conditional logistic regression was used and the adjusted odds ratio including 95% CI was reported [32]. We assessed model fit and statistical significance at the 0.05 level.

## Results

### Patient Cohort and Baseline Characteristics

A total of 177 patients who underwent surgical resection for TNM stage II colon cancer were identified from 2003 to 2018 in the Leiden University Medical Center (*Figure 1*). Of these, 40 were excluded either because they had an incomplete follow-up period of less than five years from the date of surgery, or because tissue samples were unavailable for analysis. The cohort available for matching comprised 137 patients of which 26 developed a recurrence (case) and 111 patients did not (control). Case-control matching in a 1:3 ratio, based on pT4 status and the number of sampled lymph nodes, was achieved for 25 recurrence and 75 non-recurrence patients, resulting in a total study cohort of 100 patients.

**Figure 1.**
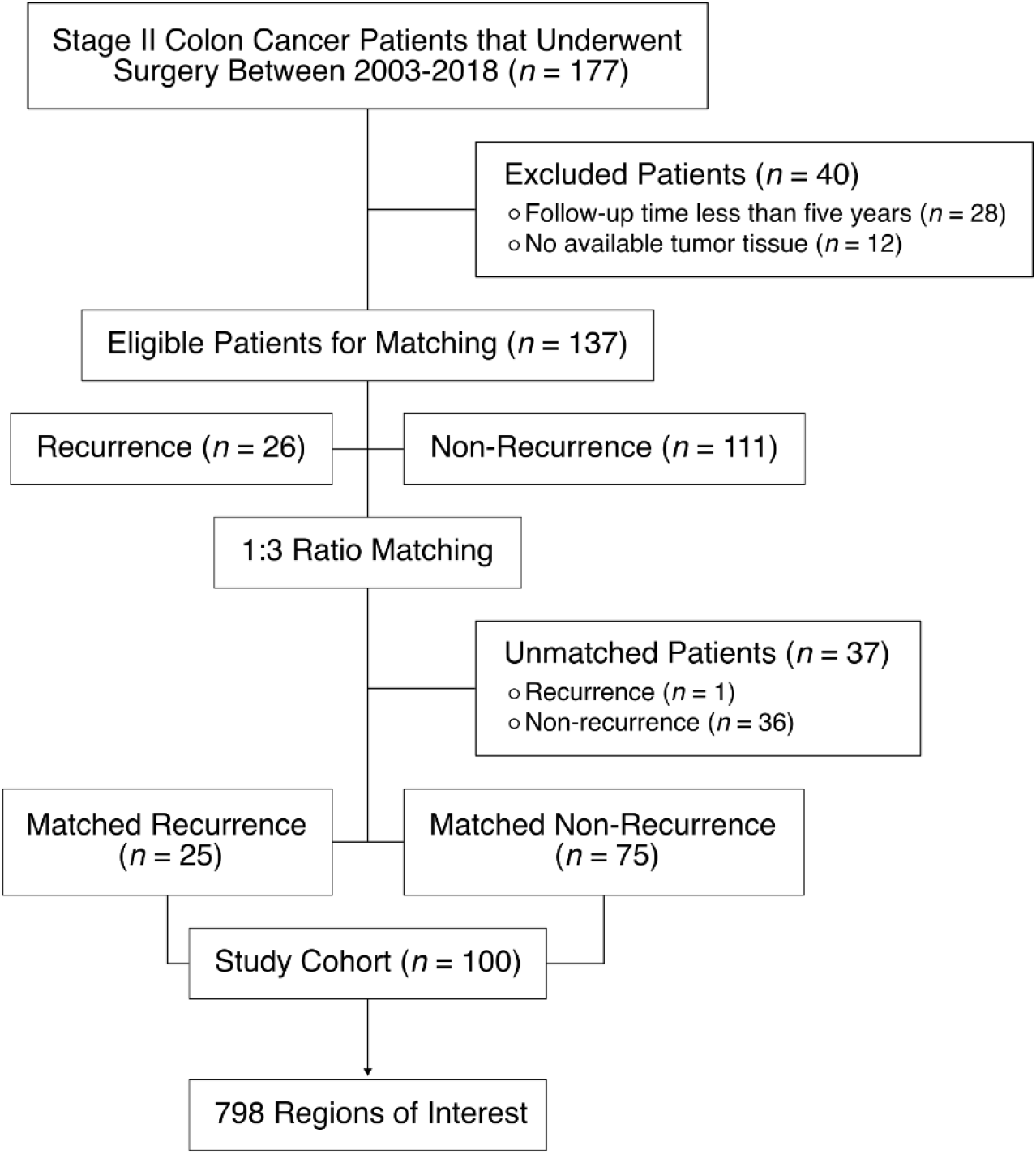
Flow diagram of the study cohort and case-control matching.

Baseline characteristics for the recurrence and non-recurrence groups are detailed in *Table 1*. Lymphovascular space invasion (LVSI) was notably more frequent in the recurrence group (40%) compared to the non-recurrence group (14.7%) (*p* = 0.016). No other patient or disease-specific characteristics showed statistically significant differences at baseline, suggesting that the matching process achieved effective balance between groups. Both groups had a median age of 69 years. In the recurrence group, four patients (16%) underwent surgery in an acute setting compared to five patients (7%) in the non-recurrence group (*p* = 0.209). The median tumor diameter was 47 mm in the recurrence group and 41 mm in the non-recurrence group (p = 0.817). Additionally, three patients (12%) in the recurrence group received adjuvant chemotherapy compared to four patients (5%) in the non-recurrence group (*p* = 0.374).

**Table 1.** Patient and tumor characteristics of the matched study cohort.

|  | Recurrence<br>(N = 25) | Non-recurrence<br>(N = 75) | p-value |
| --- | --- | --- | --- |
| Sex |  |  | 0.634<sup>‡</sup> |
| Female | 8 (32%) | 30 (40%) |  |
| Male | 17 (68%) | 45 (60%) |  |
| Age (years) |  |  | 0.527<sup>#</sup> |
| Median (IQR) | 69 (12) | 69 (17) |  |
| Clinical setting |  |  | 0.222<sup>±</sup> |
| Acute | 4 (16%) | 5 (7%) |  |
| Elective | 21 (84%) | 70 (93%) |  |
| Residual tumor (R) |  |  | NA |
| R0 | 25 (100%) | 75 (100%) |  |
| R1/R2 | 0 (0%) | 0 (0%) |  |
| Tumor diameter (mm) |  |  | 0.956<sup>#</sup> |
| Median (IQR) | 47 (25) | 41 (19) |  |
| Laterality |  |  | 0.133<sup>‡</sup> |
| Right-sided | 9 (36%) | 42 (56%) |  |
| Left-sided | 16 (64%) | 33 (44%) |  |
| Invasion depth |  |  | 1.000<sup>‡</sup> |
| pT3 | 20 (80%) | 61 (81%) |  |
| pT4 | 5 (20%) | 14 (19%) |  |
| Lymph nodes sampled |  |  | 0.463<sup>#</sup> |
| Median (IQR) | 14 (13) | 16 (11) |  |
| Histomorphology |  |  | 1.000<sup>±</sup> |
| Adenocarcinoma | 21 (84%) | 62 (83%) |  |
| Mucinous carcinoma | 4 (16%) | 13 (17%) |  |
| Differentiation |  |  | 0.732<sup>±</sup> |
| Moderate/well | 21 (84%) | 66 (88%) |  |
| Poor | 4 (16%) | 9 (12%) |  |
| Lymphovascular space invasion |  |  | **0.016<sup>‡</sup>** |
| Yes | 10 (40%) | 11 (14.7%) |  |
| No | 15 (60%) | 64 (85.3%) |  |
| Mismatch repair status |  |  | 0.619<sup>±</sup> |
| Proficient | 8 (32%) | 24 (32%) |  |
| Deficient | 1 (4%) | 9 (12%) |  |
| Not determined | 16 (64%) | 42 (56%) |  |
| Adjuvant chemotherapy |  |  | 0.362<sup>±</sup> |
| Yes | 3 (12%) | 4 (5%) |  |
| No | 22 (88%) | 71 (95%) |  |

Abbreviations: IQR, interquartile range; mm, millimeter; pT, pathological tumor status.
<sup>‡</sup>Chi-squared test
<sup>#</sup>Mann–Whitney U test
<sup>±</sup>Fisher's exact test

### ECM Detection Using Fluorescent Imaging of Picrosirius Red-Stained Tissues

To confirm that the Picrosirius red (PSR) fluorescence imaging method accurately detects ECM components, we compared its results with PSR polarization microscopy and established histological stains targeting specific ECM structures. Polarized light microscopy highlighted dense (red) and sparse (green) collagen fiber bundles, which were similarly detected in the red fluorescence channel of PSR imaging (*Figure 2A*). Herovici staining further differentiated mature and immature collagen, both of which were detected with PSR fluorescence, highlighting the method’s ability to visualize collagen structures (*Figure 2B*). Orcein staining highlighted elastic fibers, which, though not stained by Sirius red, emitted autofluorescence in the green channel (*Figure 2C*). This shows that PSR fluorescence imaging allows visualization of elastic fibers, unlike brightfield imaging. Lastly, Alcian blue staining identified glycosaminoglycans (GAGs), also detected in the red channel due to their spatial proximity to collagen (*Figure 2D*). Together, these results confirm that PSR fluorescence imaging reliably detects key ECM components, including collagen and elastic fibers.

**Figure 2.**
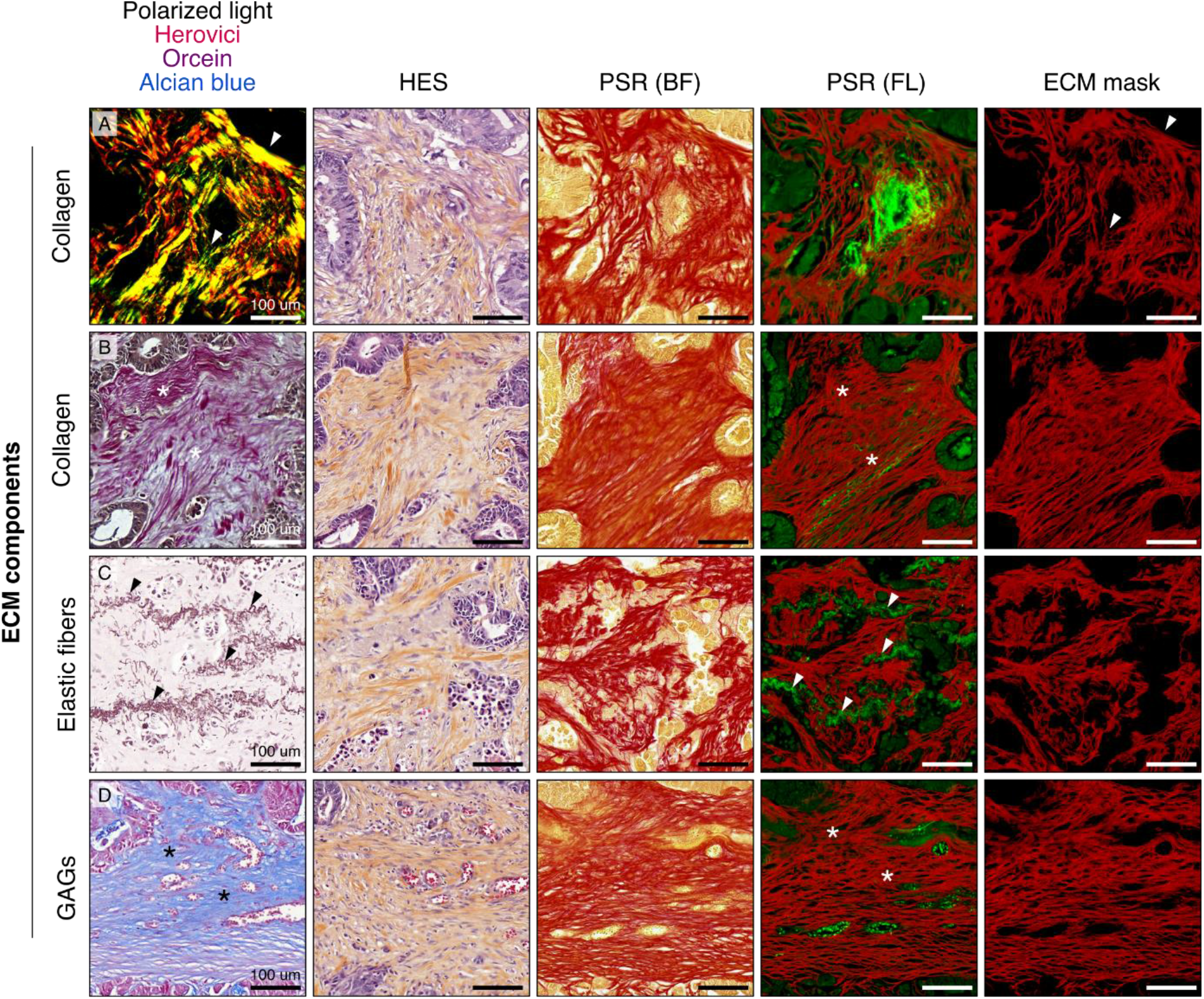
Validation of Picrosirius red (PSR) fluorescence extracellular matrix (ECM) imaging using polarized light microscopy and histochemical stainings. **(A)** Polarized light microscopy of PSR-stained tissue sections detects dense (red) and sparse (green) collagen fiber bundles (arrows). PSR fluorescence imaging detects collagen fiber bundles of varying densities, as shown in panel PSR (FL). **(B)** Herovici staining differentiates between mature (fuchsin) and immature (grey) collagen fibers, marked with asterisks. PSR fluorescence imaging detects both types of collagens in the red channel, as shown in panel PSR (FL). **(C)** Orcein staining highlights elastic fibers (indicated by arrows). These fibers are also identified in the green autofluorescence channel of Picrosirius red fluorescence imaging, as shown in panel PSR (FL). **(D)** Alcian blue staining shows the presence of glycosaminoglycans (marked with asterisks). These components are detected in the red channel of Picrosirius red fluorescence imaging, as shown in panel PSR (FL). Abbreviations: BF, brightfield; ECM, extracellular matrix; FL, fluorescence; GAGs, glycosaminoglycans; HES, hematoxylin-eosin-saffron; PSR, Picrosirius red.

### Unsupervised Clustering Identifies Three ECM Morphologies in Peritumoral Stroma

We analyzed 798 ROIs for quantitative ECM characteristics, with 224 from recurrence patients (average 8.9 ROIs per patient) and 574 from non-recurrence patients (average 7.7 ROIs per patient) (*Figure 3A*). Principal component analysis (PCA) was performed, retaining four components that explained over 80% of the total variance (*Supplementary Figure S2A-F*). Using unsupervised hierarchical clustering on these components, we identified three distinct ECM morphology clusters (*Figure 3B*) (*Supplementary Figure S3*). A detailed examination of these clusters revealed distinct ECM features in each cluster (*Figure 3C*). Cluster 1 (254 ROIs, 31.8%) was characterized by dense, aligned ECM with low porosity, exhibiting intense collagen and glycosaminoglycan (GAG) staining in Movat staining (*Figure 3D*). Cluster 2 (305 ROIs, 38.2%) featured a porous, low-density ECM configuration with prominent GAG staining and was frequently associated with substantial cellular infiltrates (*Figure 3E*). Lastly, Cluster 3 (239 ROIs, 29.9%) displayed a complex, crosslinked, and tortuous ECM structure (*Figure 3F*). These ECM morphologies were often found coexisting within individual tumors, showing notable variability in their frequency and distribution throughout the peritumoral stroma (*Figure 3G*).

**Figure 3.**
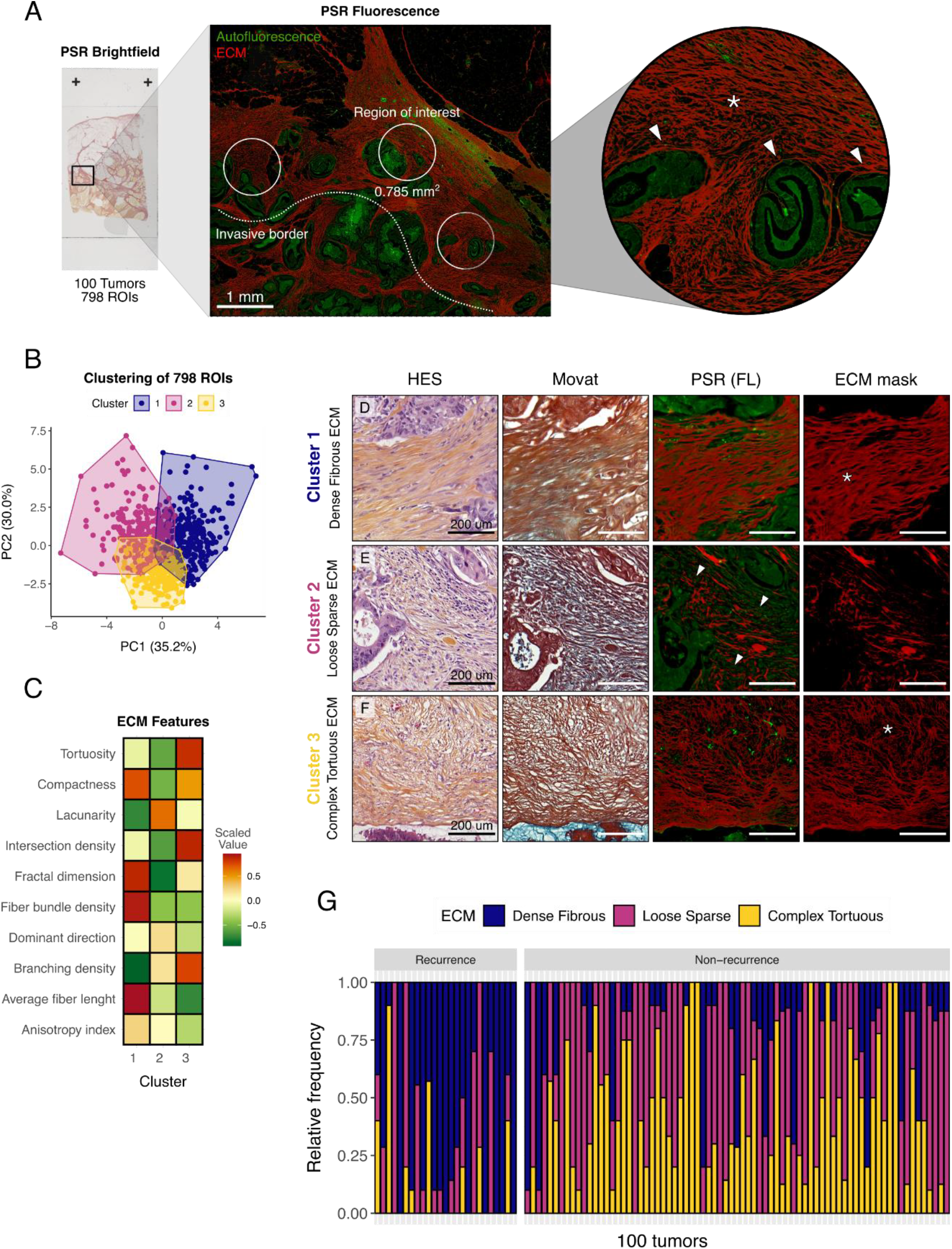
Comprehensive analysis of ECM morphology and tumor characteristics using PSR fluorescence imaging. **(A)** Example of regions of interest (ROIs) analyzed using Picrosirius red (PSR) fluorescence imaging. The asterisk marks the ECM. The arrows highlight tumor cells. **(B)** Scatterplot showing the first two principal components (PCs) from 798 ROIs, with three distinct ECM morphology clusters identified through unsupervised clustering. **(C)** Heatmap illustrating standardized ECM morphological parameters for each identified cluster to facilitate comparison. **(D)** Histological images of dense fibrous ECM (Cluster 1). The Movat panel illustrates the presence of collagen (yellow/orange hues) or glycosaminoglycans (GAG, blue hues). The asterisk highlights an area with dense ECM. **(E)** Histological images of loose sparse ECM (Cluster 2). Intense GAG staining and low collagen content were observed in these regions. Arrows indicate areas with substantial cellular infiltration. **(F)** Histological images of complex tortuous ECM (Cluster 3). The asterisk marks tortuous, crosslinked ECM. **(G)** Stacked bar chart displaying the relative frequency of invasive border ECM morphologies observed in each tumor. Abbreviations: ECM, extracellular matrix; FL, fluorescence; GAG, glycosaminoglycan; HES, hematoxylin-eosin-saffron; PC, principal component; PSR, Picrosirius red; ROI, region of interest.

### Dense Fibrous ECM is Associated with Increased Risk of Recurrence

We then studied the prevalence of ECM morphologies in primary tumors from both recurrence and non-recurrence patients. In recurrence cases, 65.6% (147) of the ROIs exhibited dense fibrous ECM, 23.2% (52) had loose sparse ECM, and 11.2% (25) showed complex tortuous ECM (*Figure 4A*). ECM density heatmaps generated from PSR fluorescence images of recurrence patients highlighted areas of dense ECM within their primary tumors (*Figure 4B*). In non-recurrence patients, 18.6% (107) of the ROIs showed dense fibrous ECM, 44.1% (253) exhibited loose sparse ECM, and 37.3% (214) displayed complex tortuous ECM (*Figure 4C*). Large regions of low-density ECM were observed in primary tumors of recurrence patients (*Figure 4D*).

**Figure 4.**
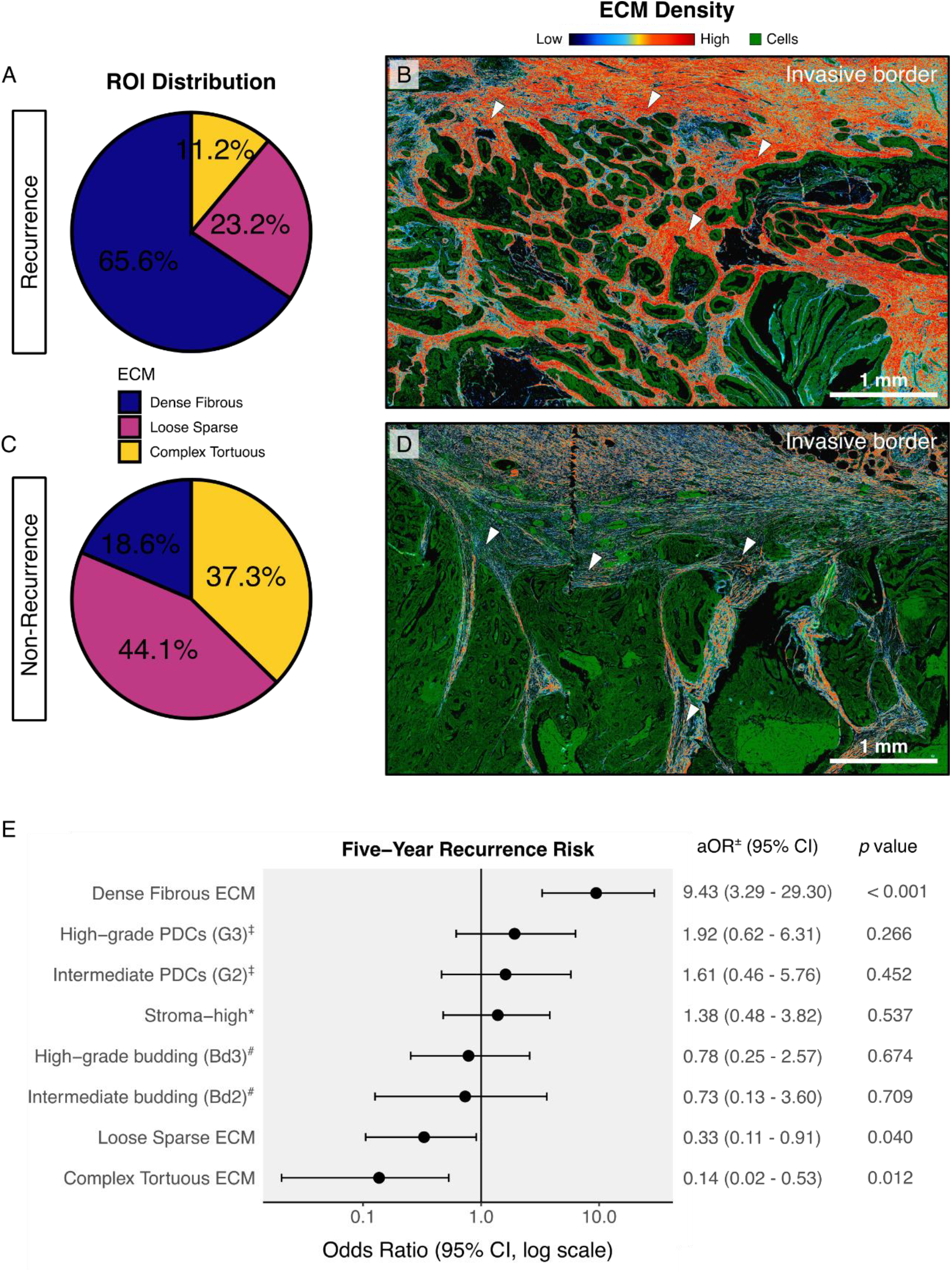
ECM morphology profiles and recurrence risk. Pie charts showing the proportion of different ECM morphology profiles in primary tumors from **(A)** recurrence patients and **(C)** non-recurrence patients. Representative ECM density heatmaps based on ECM pixel intensity from Picrosirius red fluorescence imaging, illustrating ECM density in **(B)** a recurrence patient and **(D)** a non-recurrence patient. Arrows indicate regions of high and low ECM density, respectively. **(E)** Forest plot displaying adjusted odds ratios and 95% confidence intervals for five-year recurrence risk. Predictor variables include tumors with ≥50% dense fibrous ECM, tumors with ≥50% loose sparse ECM, tumors with ≥50% complex tortuous ECM, tumor budding grade, poorly differentiated cluster grade, and tumor-stroma ratio category. Odds ratios are adjusted for matched sets and lymphovascular space invasion. Abbreviations: aOR, adjusted odds ratio; CI, confidence interval; ECM, extracellular matrix; PDC, poorly differentiated cluster. ^‡^Reference: Low-grade PDC (G1) ^*^Reference: Stroma-low #Reference: Low-grade tumor budding (Bd1) ^±^Adjusted for the matched sets and lymphovascular space invasion

To assess the link between ECM morphology and tumor recurrence, we performed conditional logistic regression, adjusted for the matched sets and LVSI, with five-year recurrence as the outcome. ROI data were aggregated into tumor-level proportions of each ECM profile, creating binary predictor variables using a 50% threshold. Tumors were categorized as having dense fibrous, loose sparse, or complex tortuous ECM if 50% or more of ROIs matched that profile. Adjusted logistic regression showed that tumors with ≥50% dense fibrous ECM had a higher recurrence risk (aOR 9.43, 95% CI 3.29-29.30, *p* < 0.001), while those with loose sparse (aOR 0.33, 95% CI 0.11-0.91, *p* = 0.040) or complex tortuous ECM (aOR 0.14, 95% CI 0.02-0.53, *p* = 0.012) had a lower risk (*Figure 4E*). Other factors, including stroma-high tumors (aOR 1.38, 95% CI 0.48-3.82, *p* = 0.537), intermediate-grade tumor budding (aOR 0.73, 95% CI 0.13-3.60, *p* = 0.709), high-grade tumor budding (aOR 0.78, 95% CI 0.25-2.57, *p* = 0.674), intermediate-grade PDCs (aOR 1.61, 95% CI 0.46-5.76, *p* = 0.452), and high-grade PDCs (aOR 1.92, 95% CI 0.62-6.31, *p* = 0.266) were not significantly associated with the recurrence.

### Association of ECM Profiles with Tumor Budding, Poorly Differentiated Clusters, and Tumor-Stroma Ratio

Tumor budding (TB) and poorly differentiated clusters (PDC) were analyzed across distinct ECM morphologies, revealing morphology-dependent variations in cellular features. TB and PDC were assessed in ECM-matched ROIs (*Figure 5A-B*). TB was significantly higher in regions with loose sparse ECM morphology compared to dense fibrous ECM (mean bud count 9.3 vs. 7.4, *p* < 0.005) and complex tortuous ECM (9.3 vs. 5.4, *p* < 0.005) (*Figure 5D*). Comparisons between dense fibrous and complex tortuous ECM showed no significant difference in TB (7.4 vs. 5.4, *p* = 0.149). PDC counts were elevated in areas with dense fibrous ECM morphology, showing significant differences compared to both loose sparse (mean 5.5 vs 3.8, *p* = 0.004) and complex tortuous ECM (5.5 vs. 4.0, *p* = 0.036), yet no significant difference was observed between loose sparse and complex tortuous ECM regions (3.8 vs. 4.0, *p* = 0.685) (*Figure 5E*).

**Figure 5.**
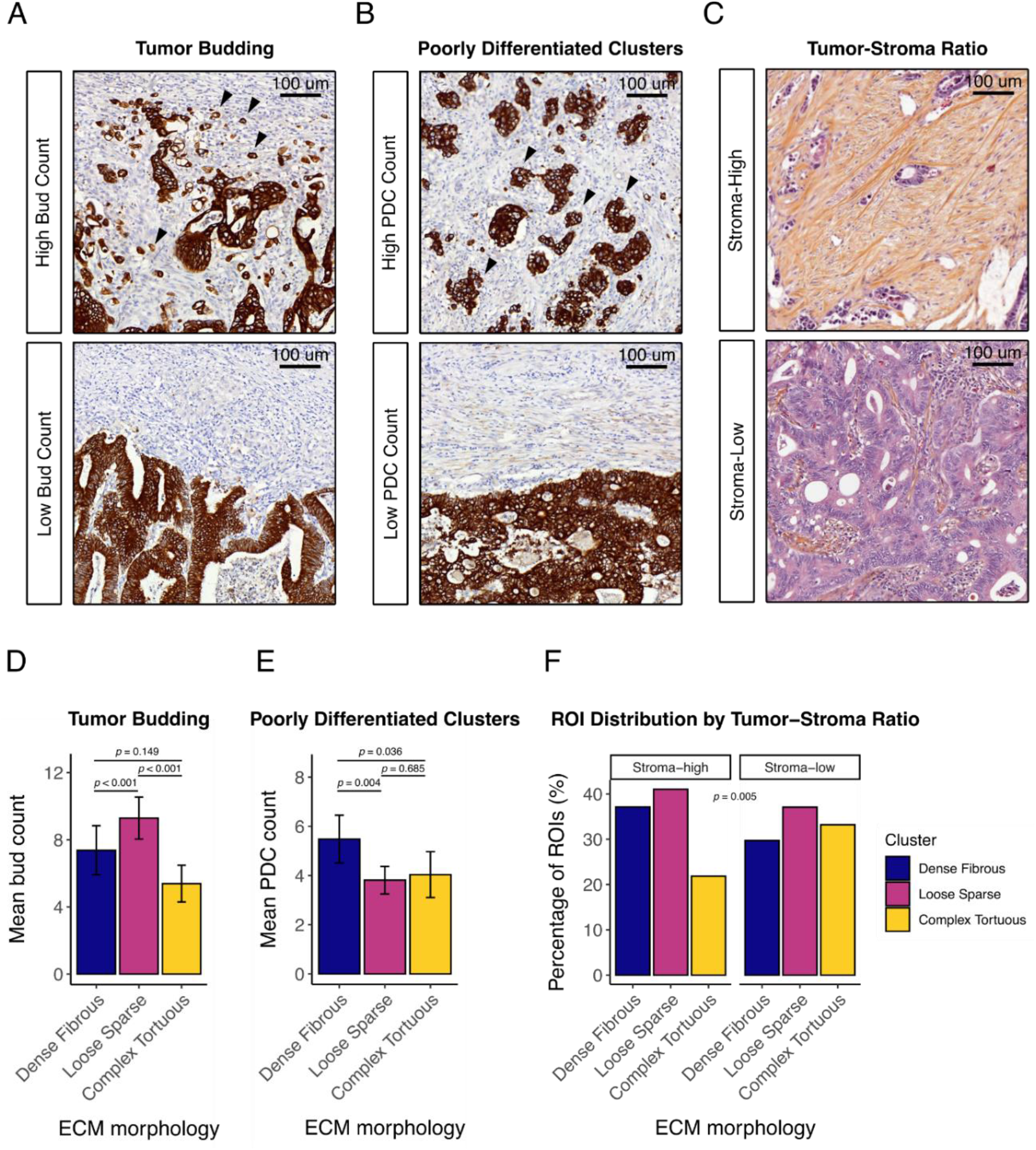
ECM morphology, tumor budding, poorly differentiated clusters, and tumor-stroma ratio. **(A)** Immunohistochemical images showing low and high tumor budding grades (arrows) stained for pan-cytokeratin. **(B)** Images showing low and high poorly differentiated cluster (PDC) grades (arrows) with pan-cytokeratin staining. **(C)** Histological images of tumors with low and high stroma ratios, stained with hematoxylin-eosin-saffron. **(D)** Bar chart of the mean tumor bud counts across ECM profiles. **(E)** Bar chart of mean PDC counts per ECM profile. **(F)** Bar chart of the proportion of regions of interest (ROIs) for each ECM profile, categorized by high-stroma (left) and low-stroma (right) tumors. Abbreviations: ECM, extracellular matrix; PDC, poorly differentiated cluster; ROI, region of interest.

Lastly, we analyzed ECM profiles relative to stromal abundance, estimated by tumor-stroma ratio (*Figure 5C*). Among 100 patients, 27% had stroma-high tumors, and 73% had stroma-low tumors. We observed marginal differences in ECM profile distribution between the stroma-high and stroma-low groups, albeit statistically significant (*p* = 0.005) (*Figure 5F*). Dense fibrous ECM was present in 37.1% (85) of ROIs in stroma-high tumors and 29.7% (169) in stroma-low tumors. Loose sparse ECM appeared in 41.0% (94) of stroma-high and 37.1% (211) of stroma-low ROIs, while complex tortuous ECM was found in 21.8% (50) of stroma-high and 33.2% (189) of stroma-low ROIs.

## Discussion

In this study, we investigated the relationship between peritumoral ECM morphology, tumor budding (TB), poorly differentiated clusters (PDC), and recurrence in TNM Stage II colon cancer, focusing on the ability of ECM morphology to predict recurrence risk. Using whole-slide ECM imaging, we identified three distinct ECM morphologies—dense fibrous, loose sparse, and complex tortuous—through quantitative analysis. Our findings revealed a significant association between these ECM morphologies and clinical tumor behavior. Dense fibrous ECM emerged as a potential marker of higher recurrence risk, independent of pT4 status, lymph node sampling count, or lymphovascular space invasion (LVSI). In contrast, loose sparse and complex tortuous ECM profiles were associated with a lower likelihood of recurrence.

The observation that dense fibrous ECM is strongly associated with recurrence aligns with previous studies linking highly organized and aligned collagen configurations to aggressive tumor phenotypes [7, 33]. The term ‘stiffness’ is commonly used to describe such rigid, dense ECM and is widely recognized as a hallmark of solid tumors [34]. In colon cancer, collagen-rich regions of the peritumoral ECM have been reported to exhibit significantly greater stiffness than the collagen-dense submucosal layer in healthy colon tissue [35]. This increased stiffness promotes stromal cell-mediated TGF-β signaling, triggering processes such as myofibroblast differentiation, epithelial-mesenchymal transition, and enhanced tumor cell motility [36-38]. Beyond facilitating tumor spread, matrix stiffening has also been linked to resistance to therapies, such as anti-VEGF monoclonal antibodies [39]. However, while ECM stiffness is commonly described in cancer, we observed the coexistence of various ECM morphologies within the same tumor. This spatial heterogeneity in ECM architecture suggests that localized ECM regulation may challenge the assumption that matrix stiffening is a universal feature of tumorigenesis.

Building on this, we recently identified three distinct ECM profiles in T1/T2 primary colon tumors: aligned dense, loose, and tortuous types [11]. These profiles mirror the ECM configurations identified in the present study, suggesting that ECM configurations might be conserved during colon cancer progression. However, loose ECM was predictive of lymph node metastasis in T1/T2 tumors, whereas dense fibrous ECM was associated with recurrence in T3/T4 tumors in this study. These findings suggest that while similar ECM morphologies appear across tumor stages, their predictive roles shift as tumors progress.

Interestingly, loose ECM in T1/T2 tumors was associated with higher numbers of cytokeratin^+^ single cells [11]. In this study, a similar pattern emerged in T3/T4 tumors, where loose sparse ECM regions showed the highest mean TB counts. Paradoxically, while loose ECM in T3/T4 tumors demonstrated high TB, it was linked to a lower risk of recurrence. Instead, dense fibrous ECM strongly predicted recurrence, with tumor regions containing this particular ECM morphology showing significantly more PDCs than other ECM morphologies. Although the exact pathogenetic role of TB and PDCs remains unclear, these features may represent different invasion modes and metastatic potentials within distinct ECM contexts. Mechanistically, in early stages (T1/T2), loose ECM may facilitate locoregional invasion of budding cells via the lymphatic system without necessarily driving systemic recurrence. By contrast, in later stages (T3/T4), dense fibrous ECM may promote the systemic spread of larger cell clusters, increasing recurrence risk. Alternatively, the presence of dense fibrous ECM could simply represent an epiphenomenon linked to tumor maturation rather than a direct contributor to invasion. If so, the association between dense ECM and recurrence may reflect the tumor’s maturity rather than an intrinsic capacity for metastasis. The observational nature of our study underscores the challenge in distinguishing causative mechanisms from tumor-associated ECM changes. Nevertheless, our findings suggest a potential ECM maturation phenomenon, where ECM becomes progressively denser and more structured as the tumor advances. The coexistence of varied ECM morphologies within the same tumor— from loose to dense types—may reflect different phases of ECM maturation, underscoring the localized regulation of this process.

Given our results, ECM profiling offers valuable insights for patient stratification and treatment planning in Stage II colon cancer, beyond conventional TNM staging. Dense fibrous ECM, in particular, could serve as a marker for patients with increased risk of recurrence in which adjuvant chemotherapy may be considered, whereas those with loose sparse or complex tortuous ECM may require more conservative follow-up regimens, reducing the risk of overtreatment in these low-risk patients. Despite the predictive value of ECM configurations for recurrence risk, it is still unclear whether patients with these high-risk characteristics benefit from adjuvant chemotherapy. This study did not explore whether specific ECM morphologies are also indicative of therapy response. Therefore, future research should investigate whether dense fibrous ECM, associated with higher recurrence risk, can also predict patient outcomes following adjuvant chemotherapy.

Although TB and PDCs are frequently associated with poor prognosis in colon cancer, we found no association between TB or PDCs and five-year recurrence risk after adjusting for high-risk factors such as pT4 status, lymph node sampling count, and LVSI. One possible explanation for this lack of association is the established link between TB, PDCs, and T status [13, 40-44]. In these studies, high-grade TB or PDC more frequently occurred in T4 tumors, which are characterized by extensive invasion into surrounding tissues. By adjusting for T4 status, the effect of TB or PDCs on recurrence may have been diminished, as T4 tumors inherently carry a higher risk of poor outcomes. In addition, the relatively small sample size may have limited the statistical power to detect significant associations between TB, PDCs, and recurrence. Nevertheless, TB and PDCs remain key histopathological features in stage II colon cancer, with strong prognostic value for disease-free survival, and are increasingly recognized in international guidelines [15, 45-47].

ECM imaging has previously yielded clinically meaningful results. For instance, tumor-associated collagen signatures (TACS) characterized by aligned collagen fibers have been linked to poor disease-specific survival in breast cancer patients [33]. Similar associations between collagen morphologies and reduced disease-free survival have been reported in colon cancer [48, 49]. These studies commonly employ non-linear optical imaging techniques, such as second harmonic generation (SHG) and two-photon excitation fluorescence (TPEF), which offer benefits like minimal sample preparation and compatibility with standard diagnostic tissue sections (e.g., H&E-stained). However, the high cost of non-linear optical imaging equipment and the lengthy imaging times have limited their broader clinical application. To address these challenges, we developed a whole-slide ECM imaging method using Picrosirius red, an established clinical stain for diagnosing fibrotic liver disease and amyloid pathologies, to capture and quantify ECM morphology across entire tumor sections [50]. Our fluorescent imaging approach, designed for compatibility with clinical workflows, enables data-driven ECM profiling and supports integration into routine pathology. By leveraging established staining techniques and quantitative analysis, this pipeline offers a feasible path for incorporating ECM characteristics into clinical decision-making.

While this study provides valuable insights, several limitations should be acknowledged. The case-control design, although useful for identifying associations, precludes causal conclusions. Additionally, our sample size may not fully capture the diversity of ECM profiles across the broader population. Further studies should explore whether distinct ECM morphologies are linked to local invasion or distant metastasis, as we did not differentiate between locoregional and distant recurrences. Moreover, we focused primarily on the peritumoral ECM, leaving intratumoral ECM characteristics unexplored. Lastly, for research purposes, we evaluated TB and PDCs using immunohistochemical staining for cytokeratin. In clinical practice, however, these histological parameters are typically assessed on H&E-stained sections. Therefore, our findings may require validation using standard H&E staining to ensure their applicability in a routine diagnostic setting.

Moving forward, future research should integrate spatial molecular profiling with ECM morphometry to identify molecular signatures associated with recurrence and to explore how ECM morphologies evolve over time. Additionally, implementing machine learning for automated ECM feature extraction could enhance the clinical utility of our findings. These advancements could open avenues for ECM-targeted therapies designed to reshape ECM architecture, potentially reducing the risk of recurrence.

In conclusion, this study highlights peritumoral ECM morphology as a predictor of recurrence in TNM Stage II colon cancer, with dense fibrous ECM strongly associated with higher recurrence risk. Our findings emphasize the importance of ECM context, demonstrating that the predictive role of ECM profiles evolves with tumor progression. By implementing whole-slide ECM imaging, we present a feasible approach for integrating ECM profiling into clinical pathology, supporting patient stratification beyond conventional staging. While further research is needed to establish causative links and explore therapeutic implications, ECM profiling may ultimately provide a path toward personalized treatment strategies aimed at minimizing recurrence.

## Abbreviations

CI: confidence interval
ECM: extracellular matrix
EMT: epithelial-mesenchymal transition
FFPE: formalin-fixed paraffin-embedded
FL: fluorescence
GAG: glycosaminoglycan
HES: hematoxylin-eosin-saffron
LVSI: lymphovascular space invasion
PCA: principal component analysis
PDC: poorly differentiated cluster
PSR: Picrosirius red
ROI: region of interest
SHG: second harmonic generation
TACS: tumor-associated collagen signatures
TB: tumor budding
TNM: tumor node metastasis
TPEF: two-photon excitation fluorescence
TSR: tumor-stroma ratio.

## Statements and Declarations

## Funding Statement

This work was supported by grants from the Bollenstreekfonds. C.J.R. is funded by an MD/PhD grant from the Leiden University Medical Center (LUMC). None of the above parties has had a role in the design of the study, collection, analysis and interpretation of data, and writing of the manuscript.

## Conflict of Interest

All authors declare that this study was conducted in the absence of any commercial or financial relationships that may be construed as a potential conflict of interest.

## Data availability statement

Inquiries concerning data requests can be directed to the corresponding author. The source code of the custom image analysis software used in this study is publicly available at https://www.github.com/cjravensbergen/MORTEX. Further data of this study are available upon reasonable request.

## Author Contribution

C.J.R., V.S.C., and W.E.M. conceptualized the study and methodology. C.J.R., V.C., and A.S.L.P.C. acquired the data. C.J.R. and H.P. performed the formal analysis and visualization. W.E.M. acquired funding for the project. C.J.R. wrote the original manuscript draft. A.S.L.P.C., J.B., J.H.F.L., R.A.E.M.T., and W.E.M. critically reviewed the manuscript.

---

**Supplementary Figure S1.**
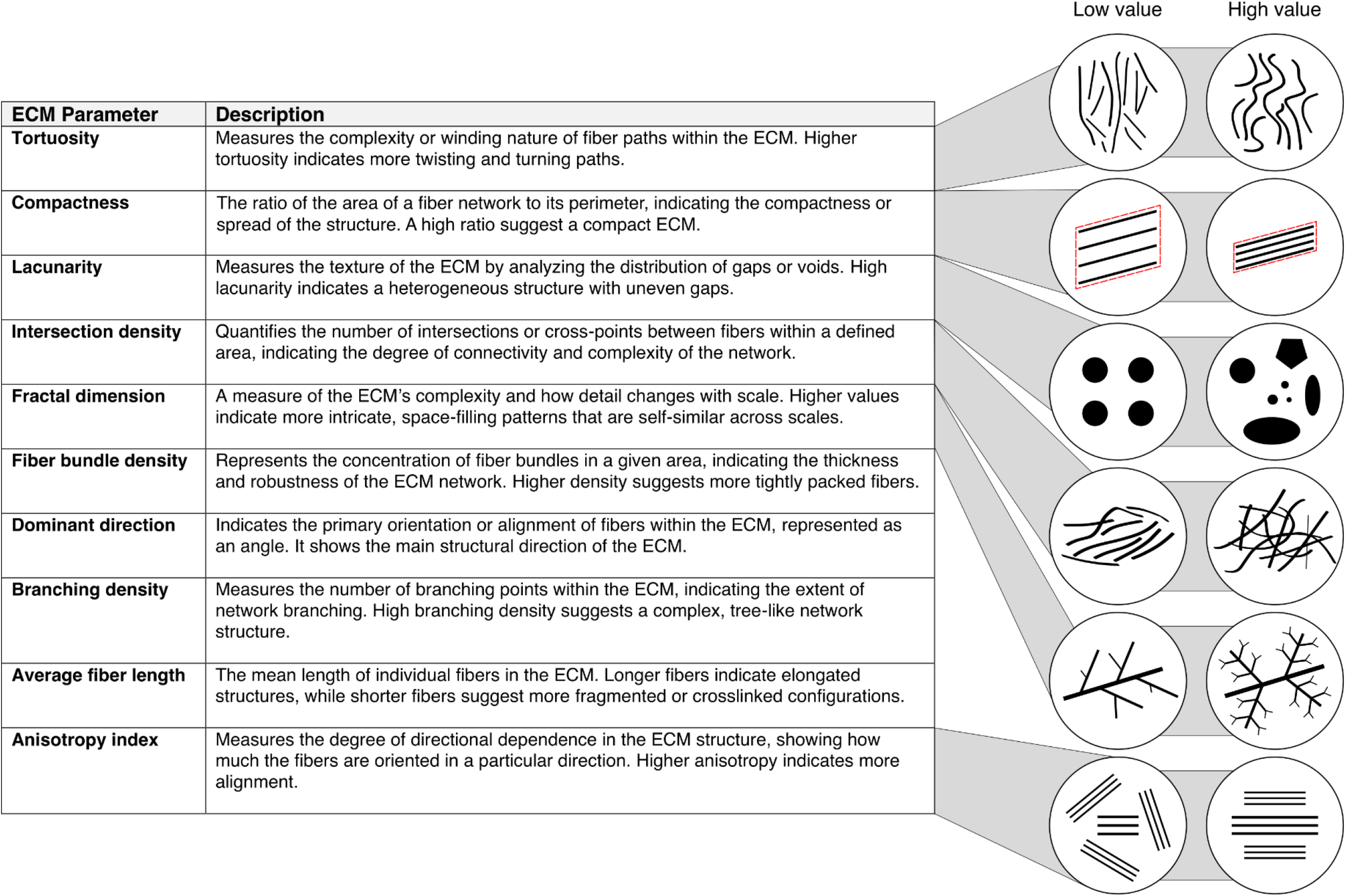
Description table of extracellular matrix (ECM) parameters. The table lists and describes the ten quantitative ECM parameters evaluated in the study. Schematic figures are provided for selected parameters to aid in understanding.

**Supplementary Figure S2.**
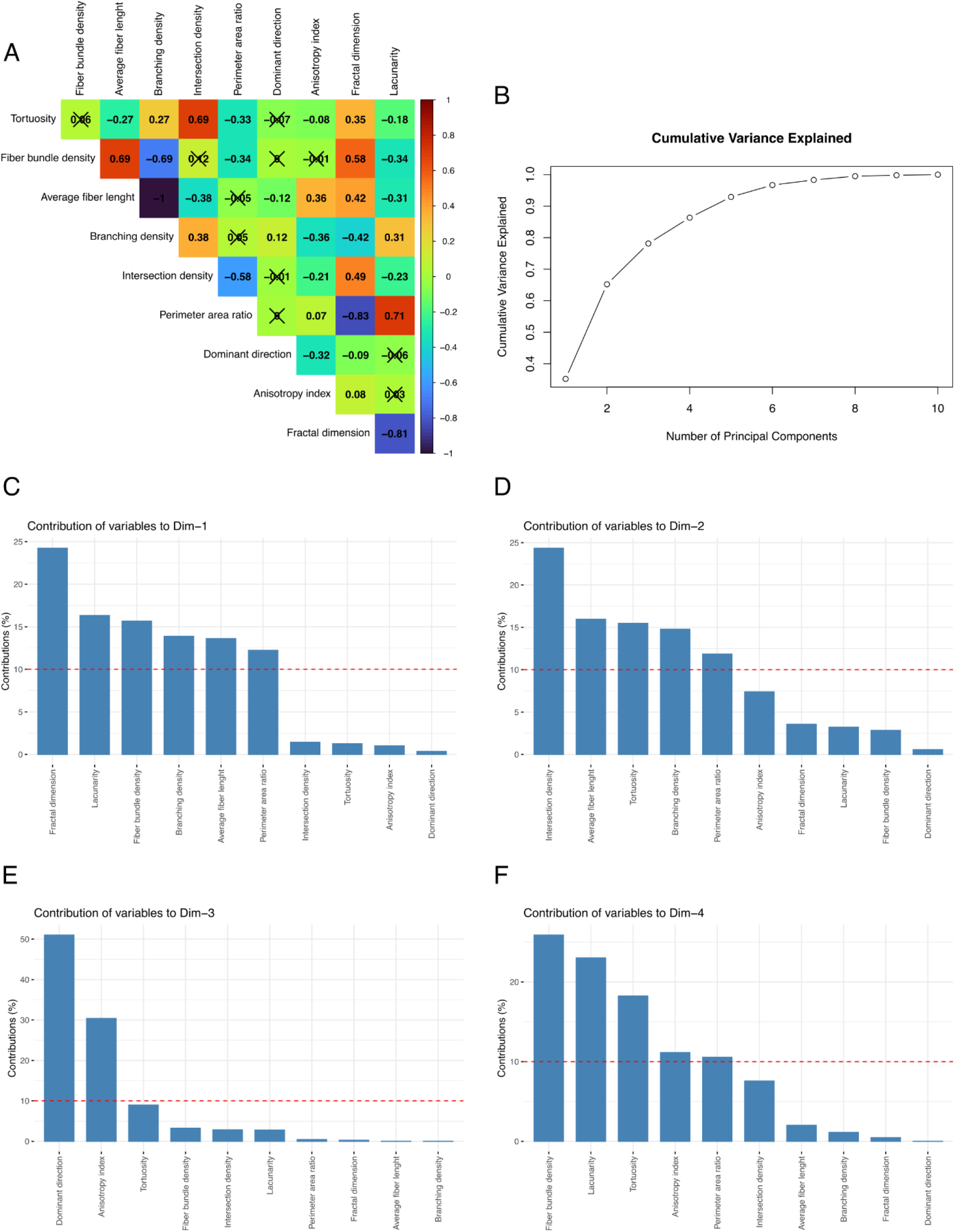
Principal component analysis of ECM parameters. **(A)** Correlation matrix showing relationships among the ten ECM parameters. **(B)** Line plot illustrating the cumulative variance explained by principal components (PCs). The plot indicates the minimum number of PCs needed to account for over 80% of the variance. **(C-F)** Contributions of individual ECM parameters to the first four principal components (PCs), highlighting their impact on each PC.

**Supplementary Figure S3.**
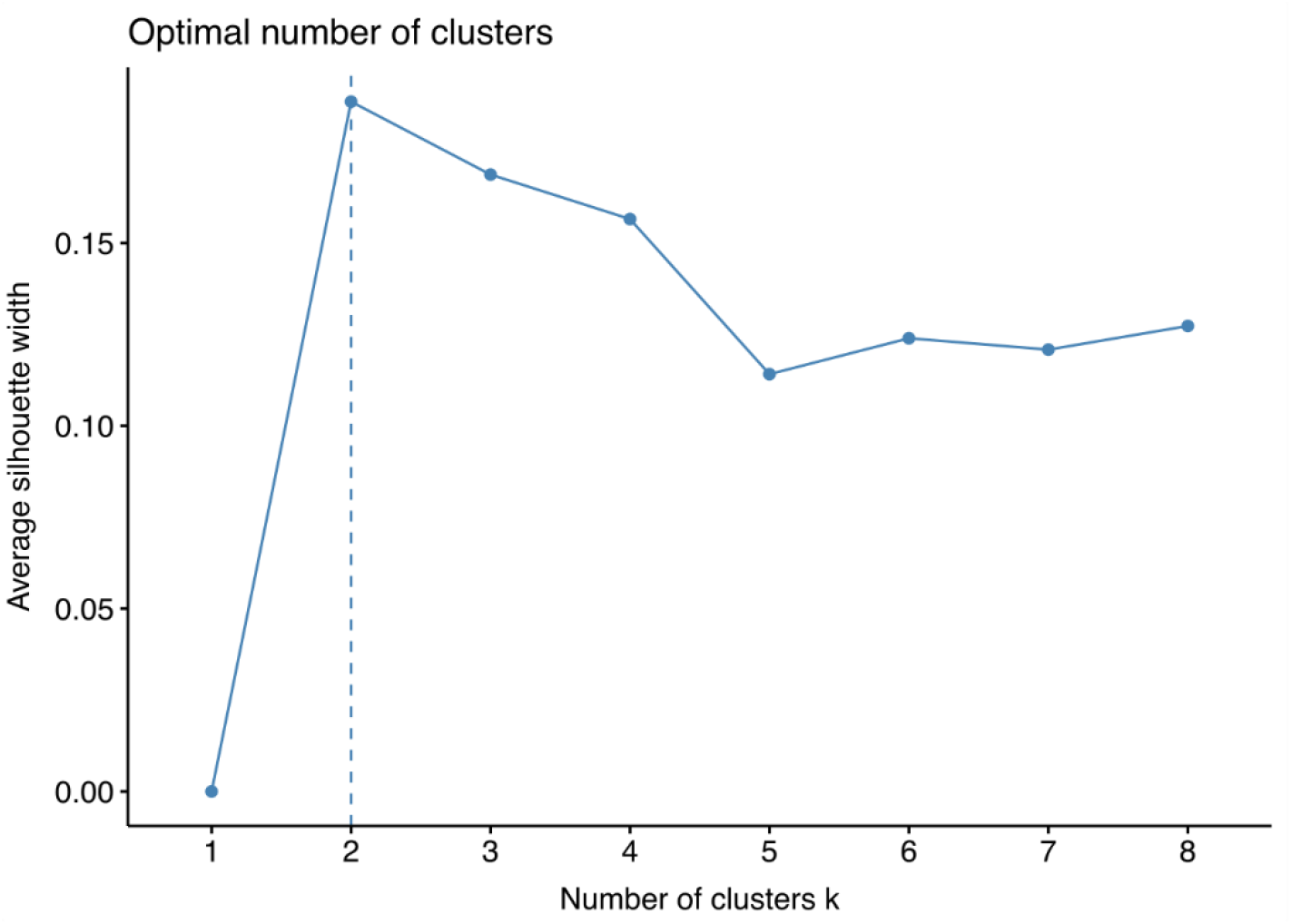
Silhouette score plot for determining the optimal number of clusters. The plot shows the silhouette scores for various cluster counts, indicating the optimal number of clusters based on the highest average silhouette score.

## Notes

https://github.com/cjravensbergen/MORTEX

### Competing Interest Statement

The authors have declared no competing interest.

